# Abscisic acid plays a key role in the regulation of date palm fruit ripening

**DOI:** 10.1101/2022.08.02.502463

**Authors:** Saar Elbar, Yochai Maytal, Isaac David, Mira Carmeli-Weissberg, Felix Shaya, Yaara Barnea-Danino, Amnon Bustan, Smadar Harpaz-Saad

## Abstract

The date palm (*Phoenix dactylifera* L.) fruit is of major importance for the nutrition of broad populations in the world’s desert strip; yet this crop is sorely understudied. Understanding the mechanisms regulating date fruit development and ripening is essential to customise this crop to the climatic change, which elaborates yield losses due to often too early occurring wet season. This study aimed to elucidate the mechanism regulating date fruit ripening. To that end, we followed the natural process of date fruit development and the effects of exogenous hormone application on fruit ripening in the elite cultivar ‘Medjool’. The results of the current study indicate that the onset of fruit ripening occurred once the seed had reached maximum dry weight. From this point until fruit harvest, the pulp endogenous abscisic acid (ABA) turned up and its levels consistently increased. The final stage in fruit ripening, the yellow-to-brown transition, was preceded by an arrest of xylem-mediated water transport into the fruit. Exogenous ABA application enhanced fruit ripening when applied just prior to the green-to-yellow fruit color transition. Repeated ABA applications hastened all fruit ripening processes, resulting in a significantly earlier harvest. The emerging pivotal role of ABA in the regulation of date fruit ripening is thoroughly discussed.

**Highlights:** The plant hormone abscisic acid (ABA) promotes date fruit ripening, as indicated by the enhanced change in fruit color, from green to yellow, and the enhanced rate of sugar accumulation.

## Introduction

*Phoenix dactylifera* L. (date palm) is a perennial dioecious monocot tree. The date palm thrives in semi-arid to arid regions under severe climatic conditions, usually considered as abiotic-stress for most plant species (Al-Mssallem *et al*., 2013). The date palm fruit is a staple food for millions of people in the Middle East, North Africa, Asia and America due to its high energy content. Moreover, it also holds potential health benefits as a result of the high nutrient content and bioactive compounds like flavonoids, tannins and other phenolic compounds (Marondedze *et al*., 2014; Lobo *et al*., 2014). However, despite of its immense nutritional, cultural, and economical significance in these regions, the date palm is an under-studied crop (Al-Khayri *et al*., 2015).

The date palm thrives in areas characterized by high temperatures and low humidity (Al-Mssallem *et al*., 2013). The prerequisites for date fruit ripening are high temperatures and no precipitations during the phase of fruit development (Lobo et al., 2014). Heat unit requirements for fruit ripening vary with cultivar and can range between 1100 to 2500 hours above 18 °C (Lobo et al., 2014). High humidity during the flowering season or at later stages of fruit development, result in various physiological disorders and, hence, also limit the area of date fruit commercial production (Yahia and Kader, 2011). In recent years, traditional date palm regions are affected by global climate changes, which too often drive earlier monsoons that have detrimental consequences on fruit quality and marketability (Haris *et al*., 2019; Hussain *et al*., 2020). In parallel, the rapid expansion of commercial date palm orchards to suboptimal climate regions has challenged completion of the process of fruit development and ripening before the late-summer temperature decline, humidity upsurge, and early rain events which result in fruit deterioration and significant yield loss (Awad, 2007). Facilitated fruit development, particularly ripening, may reduce fruit deterioration and prevent the significant yield losses caused by adverse weather events.

### Date palm fruit development

Date palm is a dioecious species. Each tree is either male or female, as defined by the chromosomal mechanism responsible for sex determination in dates (Siljak-Yakovlev et al., 1996; Al-Dous et al., 2011; Al-Mahmoud et al., 2012). In female date trees, a greenish bract, called the spathe, encloses the immature inflorescence (Fig. S1). As the spathe becomes brown, it splits longitudinally, exposing the entire inflorescence for pollination. The date palm inflorescence is composed of hundreds of flowers. The female flowers are borne on thin structures called rachillae, strands or spikelets, which themselves are borne on flat, tapering peduncles (also called rachises) originating in the axis of leaves that were developed during recent growing seasons (Fig. S1; Chao and Krueger, 2007; Salomón-Torres et al., 2021). In the agricultural practice, upon spathe fracture, the inflorescence is pollinated with pre-collected pollen (Lobo et al., 2014).

Traditionally, date fruit development is categorized into five distinct stages represented by immature creamy-white (Hababouk), immature green (Kimri), maturing yellow fruit (Khalal), soft brown (Rutab) and hard raisin-like (Tamr) fruits, respectively (Chao and Krueger, 2007). Following pollination, there is a 4-6-week phase of stagnation, during which, no growth in fruit size or biomass can be detected (Hababouk). Toward the end of this stage, one ovule in each carpellate flower develops into a fruit, whereas the other two carpels degenerate (Slavković *et al*., 2016). Then the selected ovule grows through a combination of cell divisions and cell expansions until the fruit reaches its maximum size (Kimri). During this stage, there is a continuous increase in reducing sugars while moisture content is at a peak (up to 85%; Barreveld, 1993). Once the fruit reaches maximum size, the seed is fully mature (can germinate if dispersed) and ripening of the fruit pericarp begins (Khalal). This involves gradual chlorophyll degradation and carotenoids processing, leading to pericarp color change from green to yellow alongside an increase in fruit sucrose concentrations (Barreveld, 1993; Bernstein, 2004). When the pericarp Brix value reaches about 40% (Ben-Zvi *et al*., 2017), fruit color changes from yellow to brown (Rutab), which typically starts at the distal tip and progressively continues towards the calyx end. During this phase, sucrose is converted to reducing sugars, the fruit softens and tannins precipitate, leading to astringency loss. Finally, the reduction in fruit water content yields a ripe dry fruit with a sugar content of 70-80% (Tamr), a phenomenal level that has been documented so far only in date fruit (Barreveld, 1993; Lobo et al., 2014).

### The regulation of date fruit ripening

Fruit ripening is a complex process during which various metabolic pathways are orchestrated in a timely manner. These include chlorophyll breakdown, pigment synthesis, sugar accumulation, acid degradation, aroma-related volatile production and other processes (Coombe, 1976; Kumar *et al*., 2014). Traditionally, fruit ripening is classified as climacteric vs. non-climacteric. In climacteric fruit, ripening is characterized by a distinct peak in ethylene production followed by an abrupt peak in respiration. In such fruit, ethylene functions as a key regulator, coordinately activating a wide arsenal of processes leading to fruit ripening (Klee and Clark, 2010). In non-climacteric fruits, no significant increase in respiration or ethylene production is detected during the course of fruit ripening. Instead, accumulating data exploring the mechanism underlying the regulation of non-climacteric fruit ripening suggest a key role for the plant hormone abscisic acid (ABA; Wills and Ku, 2002; Jia et al., 2011; Zaharah et al., 2012; Ferrero et al., 2018; Karppinen et al., 2018). In certain cases, ethylene is involved in the regulation of specific aspects of non-climacteric fruit ripening but not in the coordinated induction of the entire process (Stewart and Wheaton, 1972; Katz *et al*., 2004; Li *et al*., 2016; Chen *et al*., 2018; Farcuh *et al*., 2018)

In dates, there is an ongoing debate about the mechanism that regulates fruit ripening. A number of studies suggest that date fruit ripening follows a climacteric pattern. These are based on a small peak in ethylene production at early stage of date fruit ripening reported for ‘Negros’ cultivar (Serrano *et al*., 2001), an increase in ethylene production during fruit ripening reported for ‘Zahdi’, ‘Derey’, ‘Sultani’ and ‘Hillawi’ cultivars (Abdel-Latif, 1988; Abbas and Ibrahim, 1996), and additional reports demonstrating that ethylene application facilitates date fruit ripening pre- and post-harvest (Musa, 2001; Awad, 2007; Al-Quarashi and Awad, 2011; Al-Saif et al., 2017). However, other reports suggest that date fruit ripening follows a non-climacteric pattern, as the rate of ethylene production and respiration decreases during fruit ripening (Rouhani and Bassiri, 1976; Lobo *et al*., 2014), proteins required for ethylene production decrease during fruit ripening (Marondedze *et al*., 2014)and exogenous ethylene application does not enhance date fruit ripening (Rouhani and Bassiri, 1977; Aljuburi et al., 2001; Kader and Hussein, 2009). The information about the role of ABA in date fruit ripening is scarce. Awad (2007) and, more recently Shareef (2020), provided evidence associating an increase in endogenous ABA concentrations with fruit ripening in the cultivars Helali and Hillawy (Awad, 2007; Shareef and Al-Khayri, 2020). However, both studies focused on the stage of fruit pericarp transition from yellow-to-brown (from ‘Khalal’ to ‘Rutab’) the final stage in fruit ripening; there have been no reports regarding earlier ripening processes. Here we aimed to investigate the mechanism regulating date fruit ripening and specifically, to study the role of ethylene and ABA in this process.

## Results

### Fruit development and ripening of ‘Medjool’ date fruit

In order to identify components regulating date fruit ripening, the natural process of ‘Medjool’ date fruit development was monitored from pollination until fruit ripening, towards harvest. During the first six weeks post-pollination (WPP), no substantial changes in fruit size or shape were observed (Fig. 1). However, starting from 6 WPP, extensive fruit growth was detected, reaching maximum fresh weight at about 17 WPP (Figs. 1A, B and S3A, B). Then at 22 WPP, following 4 weeks (17-21 WPP) during which no significant changes were observed, fruit fresh weight decreased until harvest (Figs. 1B and S3B). The reduction in fruit weight during the last phase of fruit development, corresponded with a decrease in fruit water content (WC), which began at 15 WPP and was significantly accelerated from 21 WPP until the fruit was ready to harvest (Fig. 1C). Sugar accumulation in the fruit, measured as the pericarp total soluble solids (TSS) ranged between 10% and 15% until week 16, after which a significant increase was detected, gradually rising to about 35% by 21 WPP, and then sharply shooting up to about 70% in the ready-to-harvest, fully ripe fruit (Fig. 1C). The chlorophyll fluorescence index revealed a continuous reduction in fruit-exocarp chlorophyll levels, which began at 10-12 WPP (Figs. 1D and S3C), reaching a minimum value of ∼0.4 fluorescence index units (SFR_G) by 19 WPP, which paralleled the complete fruit pericarp color transition from green to yellow (Figs. 1A, D and S3A, C). In addition, chromameter measurements of the fruit L*, a*, b* color space revealed a distinct decline in the green color, as indicated by the increase of a* between 17 WPP and 22 WPP, accompanied by an increase in the yellow color component indicated by b*, which peaked in 19 WPP, remained stable until 21 WPP and then gradually decreased. Color brightness (L*) remained stable until 21 WPP, and then declined in association with fruit pericarp color transition from yellow to brown (Fig. 1A, E). Similar, but much stronger trends were also observed in the following year (Fig. S3A, D).

**Figure 1.**
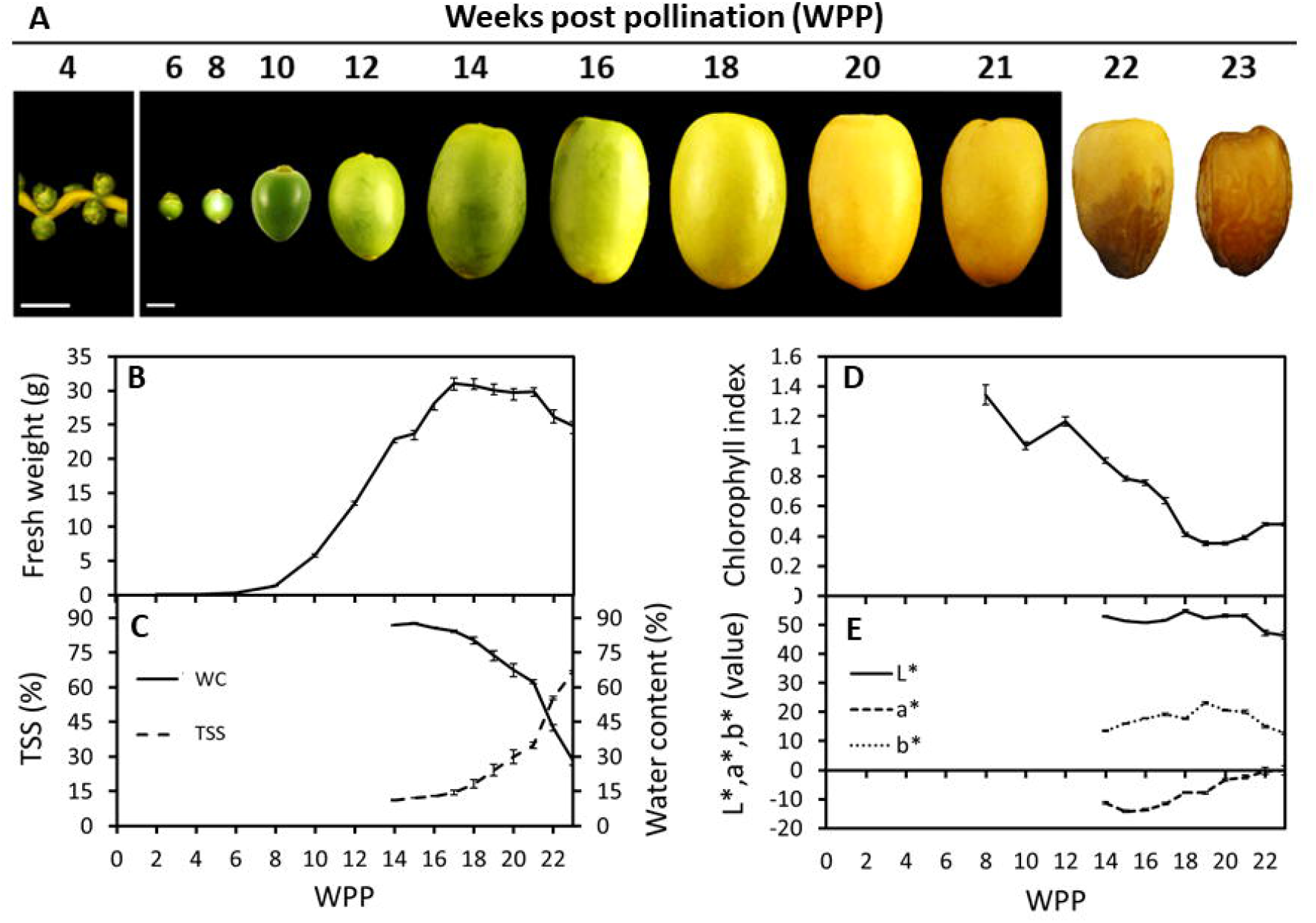
Date palm fruit development. Representative fruits were sampled and photographed along the course of fruit development (Grofit 2017). Numbers indicate weeks post-pollination (WPP; A). Fruit development was characterized through quantification of (B) fruit fresh-weight, (C) total soluble solids % (TSS) and water content % (WC), (D) chlorophyll relative fluorescence and (E) L*a*b* color space parameters. Error bars indicate standard error. Scale bar: 1 cm. The number of fruits measured at each time point was (B, D, E) n=18 or (C) n=6.

Monitoring of seed development dynamics during the course of ‘Medjool’ date fruit pericarp development, provided insight into the interrelationship between these two organs. Seed coat color gradually turned brown from 17 WPP until 21 WPP, when it was completely brown (Fig. 2A). Seed growth, as measured by dry weight, continued until 16 WPP, when the seed reached its final biomass (Fig. 2B). Seed WC decreased consistently from 13 WPP until 21 WPP and then sharply declined until harvest (Fig. 2B). In contrast to the seed, pericarp growth in dry weight recommenced at 15 WPP and gradually increased until 22 WPP, while its WC increased until 16 WPP, remained stable for about two weeks, declined slowly until 21 WPP, and then dropped sharply until fruit harvest (Fig. 2C). Note that seed biomass accumulation was completed by 16 WPP, shortly before the fruit reached its maximum size, and prior to fruit pericarp color change from green to yellow that occurred at 17-18 WPP (Figs. 1 and S3).

**Figure 2.**
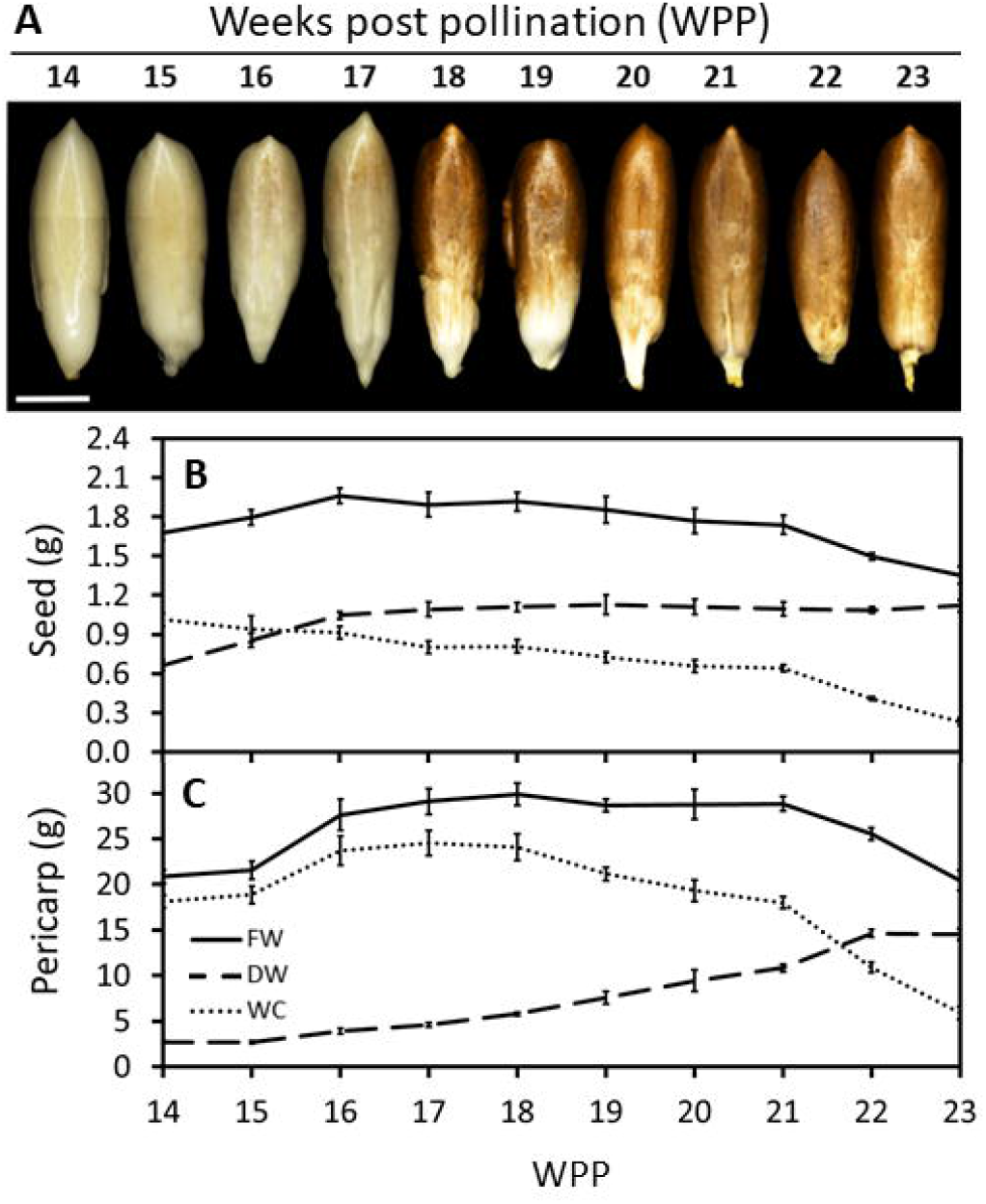
Date palm seed and fruit development. Representative fruits were sampled and seeds were separated from the pericarp tissue (Grofit 2017). (A) Seeds were photographed at different stages in fruit development as indicated by number of weeks post pollination (WPP). Dry weight (DW) and water content (WC) were quantified for both (B) seed and (C) pericarp. Note the substantial reduction in WC, in both seed and fruit pericarp, starting from 21 WPP. Error bars indicate standard error. Scale bar: 1 cm. The number of fruit measured at each time point was n=6.

Altogether, the data accumulated over the two seasons, imply two major regulatory junctures. The first, as the seed biomass reaches the maximum weight value, about 1-2 weeks before the pericarp tissues reach the maximum fresh weight (Fig. 2). This stage seems to coincide with the induction of fruit ripening, associated with the fruit pericarp color transition from green-to-yellow. The second, is just prior to the final stage in date fruit development, when a sharp reduction in fruit water content is coupled with a sharp increase in pericarp sugar levels. At this point, these changes coincide with the final color conversion of the fruit pericarp from yellow-to-brown, as the fruit becomes ready to harvest.

### Water transport to the fruit pericarp is interrupted prior to the yellow-to-brown transition

In order to understand the processes regulating the dynamics of fruit water status, the xylem flow was followed at different stages in fruit development using the water-soluble, xylem-transportable dye safranin-O. Fruit-carrying strands were detached at different stages along the course of fruit development, placed in a safranin O solution, and then examined for dye appearance in both the strand vascular system and the fruit pericarp (Figs. 3, S4 and S5). Soon after pollination, the dye was observed in all female flowers examined, and reached either all three independent ovaries of the female flower or only one of the three (Fig. S4). During the date fruitlet abscission period (5 WPP), the staining method easily discriminated between fruitlets prone to abscise and those due to persist (Fig. S5). In fruitlets that were prone to abscise, which detached from the strand in response to very slight mechanical caress, the dye streaming that was detected in the strand’s vascular system avoided the fruitlet tissues. In contrast, no interruption of the dye stream was observed in the persisting, strongly-attached fruitlets. From 6 WPP to 18 WPP, xylem transport to the fruit was not interrupted and the dye was observed in the vascular system of both the strand and the fruit pericarp (Fig. 3 and Fig. S6). Yet, at 20 WPP, while the strand’s vascular system was stained in all samples, the pericarp was stained only in half of the fruits examined (Fig. 3 and Fig. S6). Later on, from 22 WPP until harvest, the dye was observed only in the strand and pedicel vascular system but not in the pericarp (Fig. 3 and Fig. S6). Of note, the timing of the vascular continuity disruption, at 20-21 WPP, coincided with the sharp decline in fruit and seed WC (Figs. 1, 2 and S3) and the sharp increase in pericarp sugar content (Fig. 1C) and dry weight (Fig. 2C). To summarize, these results suggest that prior to the yellow-to-brown fruit color change, xylem-mediated water transport from the strand to the fruit is interrupted as part of the natural process of ‘Medjool’ date fruit development.

**Figure 3.**
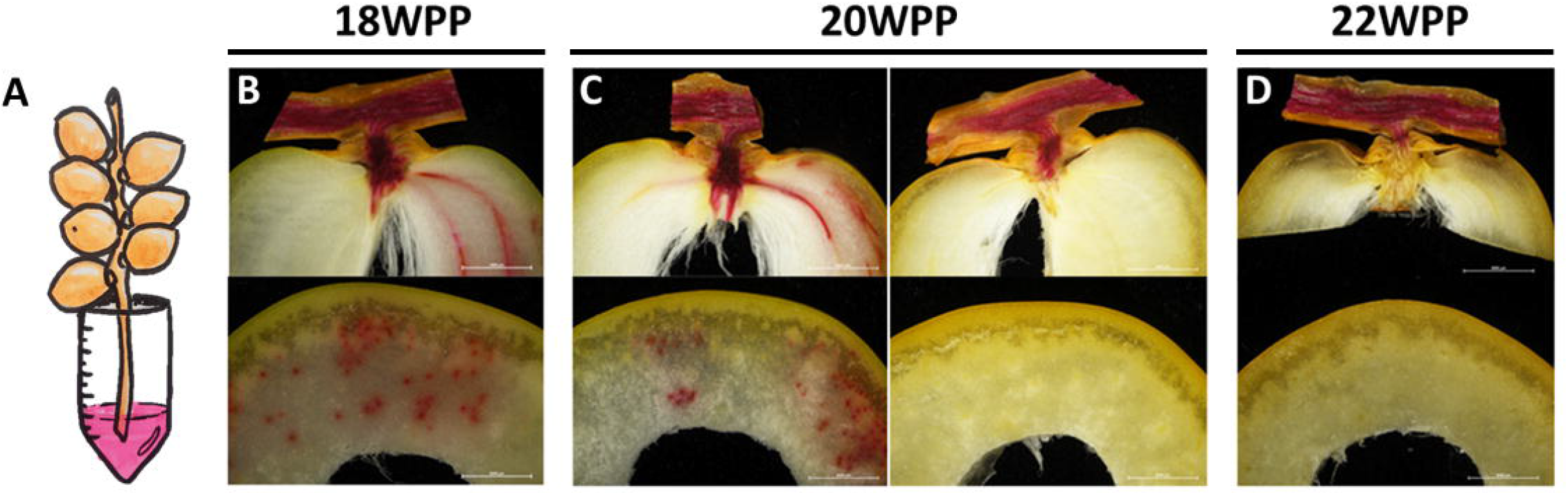
Vascular water flux arrest from the strand to the developing date fruit. Representative fruit-bearing strands were sampled along the course of fruit development (Grofit 2017) and movement of the dye Safranin O to the fruit was monitored (A) at different stages in fruit development using cross and longitudinal sections prepared (B) 18, (C) 20, and (D) 22 weeks post-pollination (WPP). Scale bar: 0.5 cm. Number of sampled strands at each time-point was n=3, each bearing 6-15 dates.

### Endogenous ABA levels increase during date palm fruit development

Endogenous ABA in the fruit pericarp was undetectable until 16 WPP. Conversely, significant endogenous ABA levels were detected starting from 18 WPP at the onset of fruit ripening, as indicated by the green-to-yellow transition (the ‘Khalal’ stage), and continuously increased throughout the process of fruit ripening until fruit harvest (Fig. 4). The rise in pericarp ABA in parallel to the process of fruit maturation suggests a regulatory role for this plant hormone in the ripening process of ‘Medjool’ date fruit.

**Figure 4.**
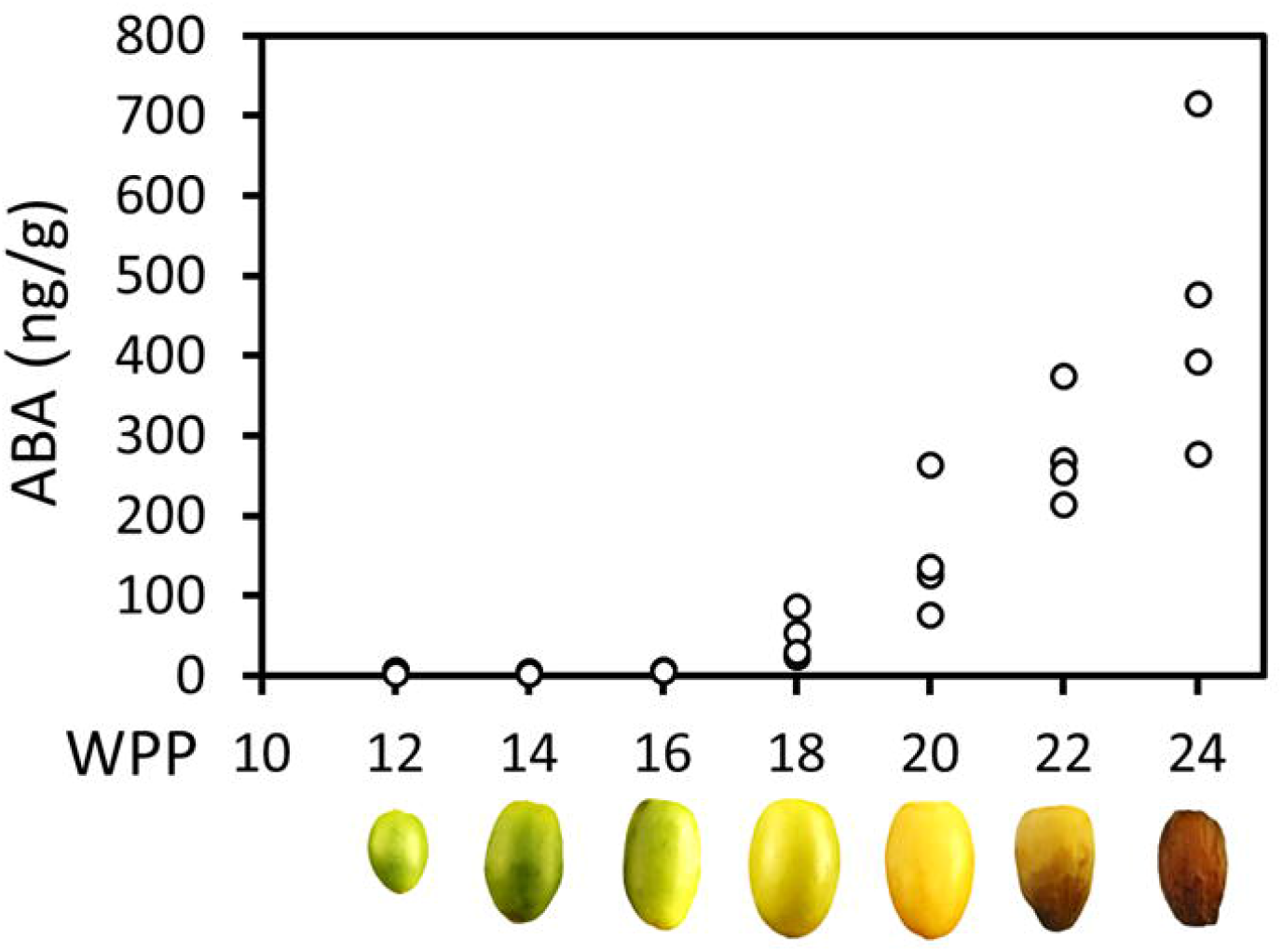
Endogenous ABA level increase throughout date palm fruit ripening. Endogenous ABA levels in representative fruits were quantified along the course of fruit development (Grofit 2017), using LC-MS. At each time point, four independent fruits from each treatment group served as biological replicates. ABA content was normalized to fresh weight.

### Exogenous ABA application facilitates fruit ripening

To better understand the role of ABA, ethylene, or a combination of the two, in date fruit ripening, individual fruiting strands were treated in the orchard with the selected hormone at 14, 16, or 20 WPP. When treated at 14 WPP, prior to a visible green-to-yellow change in fruit exocarp color, the combined ABA + ethylene treatment triggered the most prominent affect, as it facilitated pericarp color transition from green to yellow and led to a transient increase in TSS levels (Figs. S7 and S8). When applied at 16 WPP, both ABA and the combined ABA + ethylene had similar effect in initiating fruit maturation, manifested by significant green-to-yellow color changes, and an increase in TSS levels as compared to ethylene and control treatments (Fig. 5 and Fig. S9). Nevertheless, the impact of the exogenously applied ABA or ABA + ethylene at 14 and 16 WPP was transient, lasting for only 1-2 weeks after treatment (Figs S8 and S9). When applied at 20 WPP, none of the hormone treatments affect fruit development, as most of the fruit had already launched their natural maturation processes (Fig. S10).

**Figure 5.**
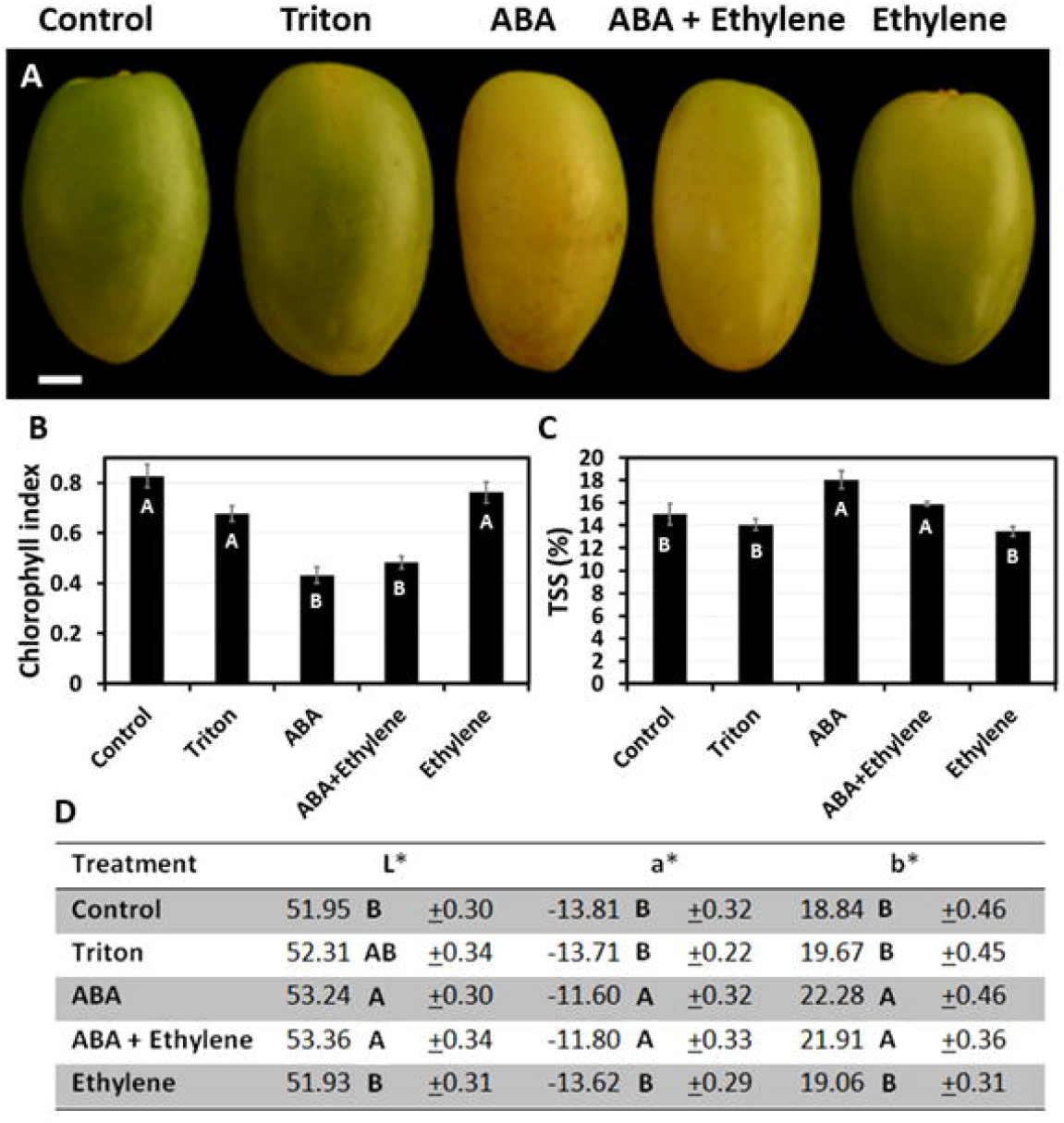
The effect of exogenous ABA and ethylene treatment applied at 16 WPP, on date fruit development. Date fruits were treated with a single treatment of ABA (ProTone™), ethylene (Ethrel®) or a combination of both, at 16 WPP on the tree (Grofit 2017). (A) Representative fruits from each treatment group were sampled and photographed one week after the treatment. Fruit ripening was characterized through quantification of (B) chlorophyll relative fluorescence, (C) total soluble solids % (TSS), and (D) color index. Error bars and “±” indicate standard error. Letters represent Tukey-Kramer multiple comparison test (B; p-value < 0.0001, C, D; p-value < 0.05). Scale bar: 1 cm. Number of fruits measured per treatment at each time-point: (B and D) n=12, or (C) n=6.

Repetitive exposure (4 applications over 13 days) of whole clusters to the ABA treatment, starting on 17 WPP, when the green fruit is at its maximum size and before the green-to-yellow transition, led to yellowing of all fruits, while all control fruits remained green (Fig. 6A, B, and D). In addition, TSS levels of all ABA-treated fruits were significantly higher compared to those of control fruits (Fig. 6C).

**Figure 6.**
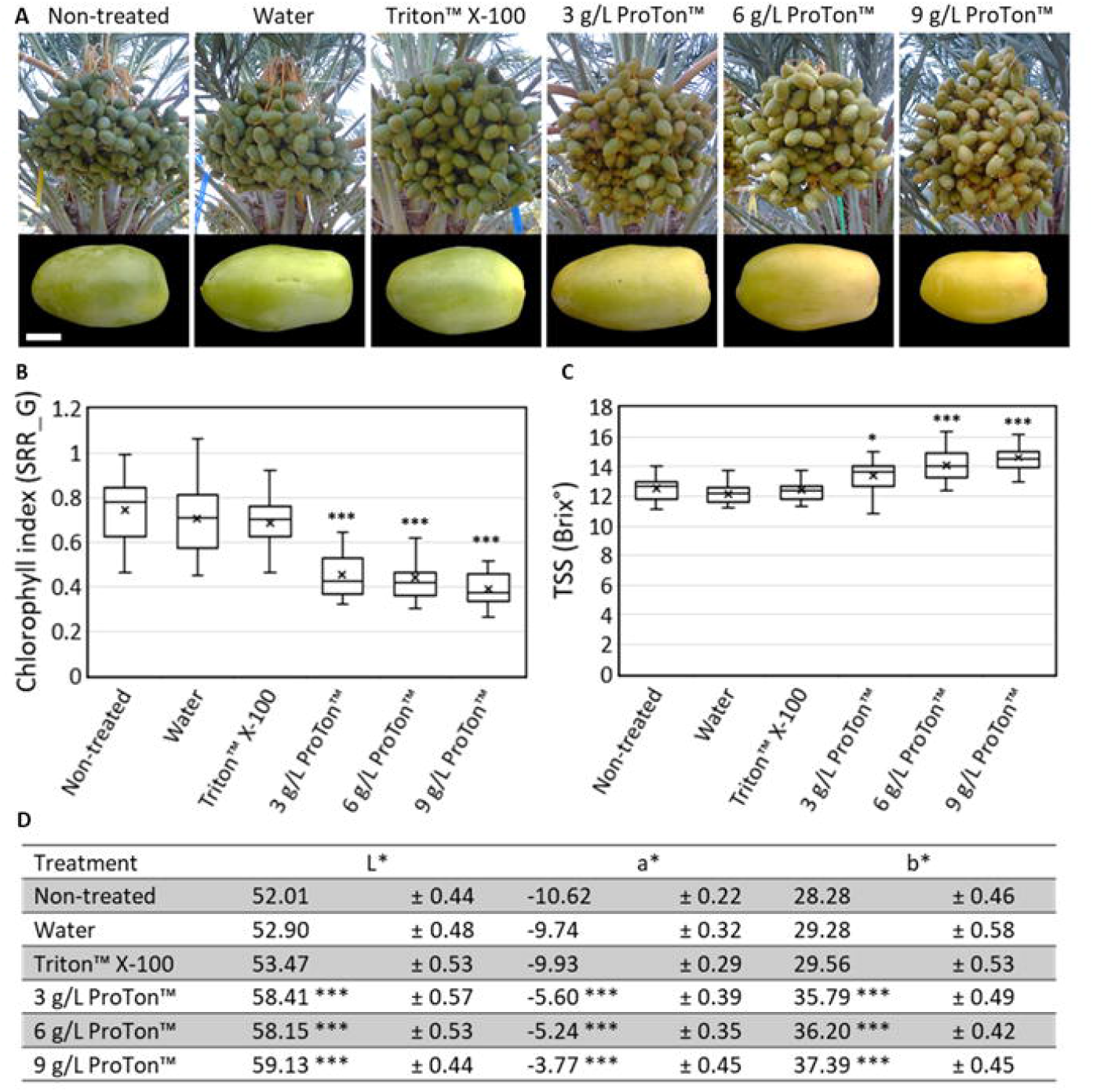
Effects of exogenous ABA treatment on date fruit maturation. Whole date fruit clusters were treated every three days (four treatments in total) with 3, 6 or 9 grL^-1^ ProTone™ (ABA), starting at 17 WPP, on the tree (Grofit 2020; A; upper panel). Representative fruits per treatment were sampled and photographed one week after the last treatment (A; lower panel). Fruit ripening was characterized through quantification of (B) chlorophyll relative fluorescence, (C) total soluble solids (TSS) (D) and the L*a*b* color space parameters. Error bars and “±” indicate standard error. All data were analyzed using Tukey-Kramer multiple comparison test (B, C, D); * indicates p-value < 0.5; *** indicates p-value < 0.0001. The number of fruits measured per treatment at each time point was n=30.

Harvest of this experiment, Grofit 2020, began at 25 WPP, and was done in 5 rounds. Total fruit yield was in the range of 6.1-8.5 kg cluster^-1^, with some tendency for higher yields in the ABA-treated clusters (Fig. S11A). While the number of fruits per cluster substantially fluctuated across treatments (Fig. S11B), mean fruit weight at harvest was significantly greater in samples treated with the higher ABA levels (6 and 9 g L^-1^ ProTone™), as compared to control and water-applied fruit (26.8 g fruit^-1^ vs. ∼22.6 g fruit^-1^ and 22.2 g fruit^-1^ for non-treated and water-treated control; Fig. S11C). Interestingly, fruit treated with a surfactant (Triton-X100) or with the lower ABA level (3 g L^-1^ ProTone™) displayed intermediate values regarding fruit weights (24.5 g and 25.4 g, respectively). When compared to controls, a much greater proportions of the ABA-treated fruit yield was harvestable in the first round (20-32%, compared to 8-15% of the total yield, respectively). On treatment with one of the two higher ABA concentrations tested, more than 60% of the fruit were harvested in the two earlier rounds, while in the non-treated and surfactant-treated control, 60-65% of the fruit were harvested in the 3^rd^ and 4^th^ rounds. Harvest from clusters treated with water, surfactant or a low ABA dose distributed quite impartially, with about 80% of the fruit collected in rounds 2 to 4. In the fifth round of harvest, only residues were left (up to 5% of the total yield), with no significant differences between treatment groups (Fig. 7C). Thus, repetitive ABA treatments accelerates date fruit ripening and precede fruit harvest.

**Figure 7.**
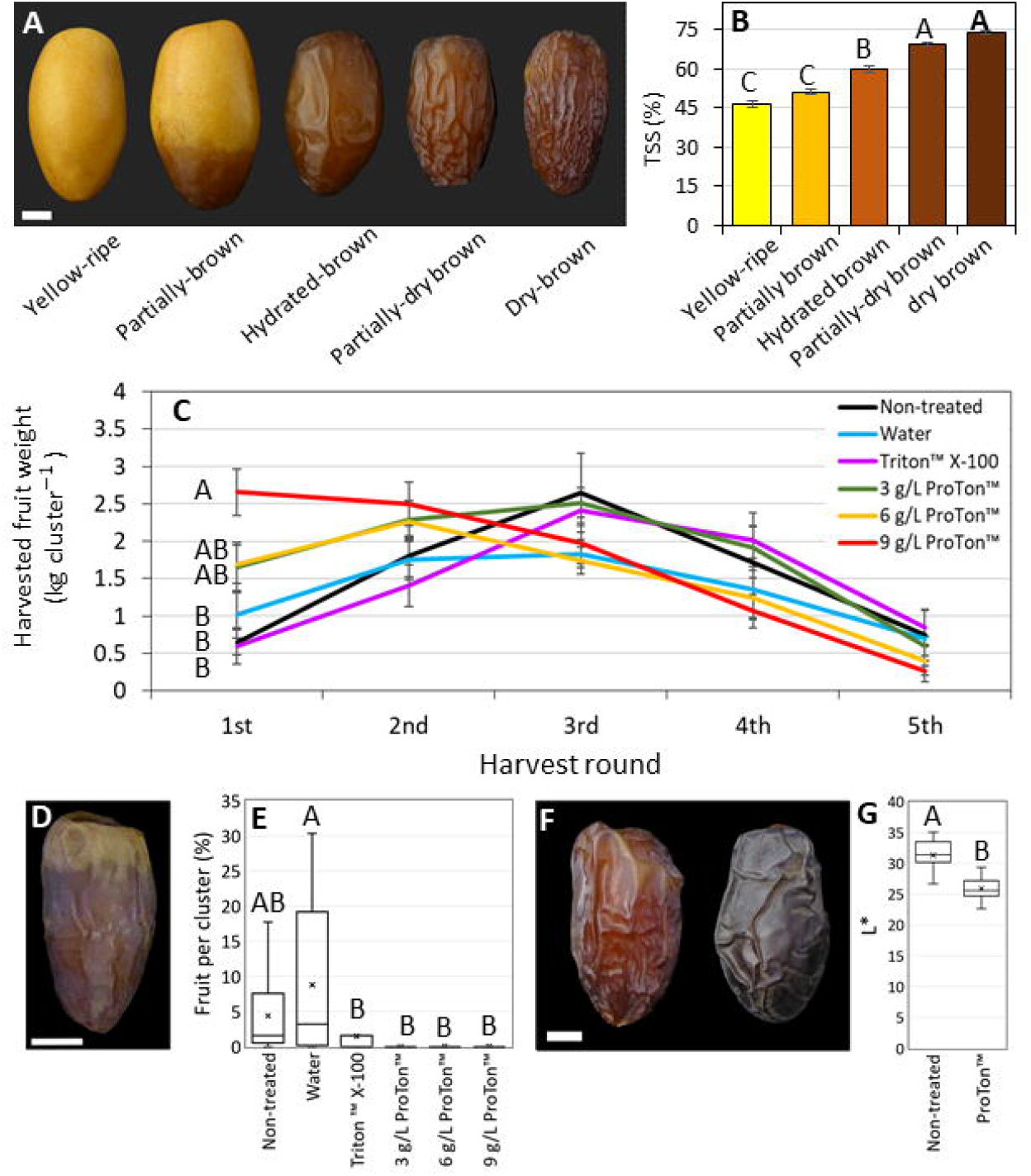
Pre-harvest ABA treatments promote fruit harvest and result in darker fruit. In the season of 2020, whole date fruit clusters were treated pre-harvest every three days (four treatments in total) with ABA (ProTone™; 3, 6 or 9 g/L), and Triton™ X-100, starting from 17 WPP. At harvest, five ripening classes were defined by (A) appearance and (B) total soluble solids % (TSS). (C) The weight of the fruit harvested from the treated clusters at each of the five harvest rounds (one round per week), is shown. (D) Typically, the first harvested fruits of this very arid habitat, are not marketable since they are too dry and do not ripen properly. (E) The percentage of such nonmarketable fruits in the first harvest round of the season was calculated. (F) Fruits treated with ABA appeared darker then the non-treated fruits. (G) Representative fruits per treatment were sampled 77 days after treatment, photographed, and measured for L* parameter of the L*a*b* color space. Error bars indicate standard error (B, C). X mark indicates mean (E, G). Data was analyzed using Tukey-Kramer multiple comparison test (B; p-value < 0.0005, C and E; p-value < 0.05, G; p-value < 0.0001). The number of fruits measured for each class was n=6 (B). The number of clusters harvested per treatment at each time-point was n=10 (C, E). The number of fruits measured for each treatment was n=30 (G). Scale bar: 1 cm (A, D, F).

In addition, it is noteworthy that the fraction of yellow fruit that easily abscised at harvest was much greater among the clusters treated with repeat ABA doses, reaching 20–30% of the fruit, compared to about 3% in the controls (Figs S12). Moreover, the fraction of partially brown fruit was higher among the control groups (14–18%), as compare to ABA-applied fruits (10-11%). Especially notable was the scarcity (less than 1%), of partially dry and dry brown fruit fractions (hydrated and dry ‘Tamr’) and the absolute dominance of hydrated brown fruit (‘Rutab’) in the ABA-treated samples, as compared to 8-15% in the controls (Fig. S12), suggesting that ABA may also affect the last stage of fruit dehydration. Subsequently, visual and spectral differences in fruit appearance were apparent; the surface of ABA-applied fruits was considerably smoother and darker compared to control fruit (Fig. 7F-7G). Lastly, repetitive ABA treatments, like the surfactant alone, reduced the fraction of prematurely-dry fruit in comparison to the non-treated or water-treated controls (<2% vs. 10-20%; Figs. 7E and S12). Taken together, repetitive ABA treatments can accelerate date fruit ripening.

## Discussion

The current study, aimed at characterizing the regulatory role of ABA in date fruit ripening, identified gradual increase in endogenous ABA levels during natural fruit ripening. One-time exogenous ABA applications accelerated the process in a transient manner, while repetitive ABA treatments enhanced fruit ripening and resulted in early fruit harvest. Interestingly, ethylene treatment did not promote fruit ripening, unless applied at very early stages in fruit development together with ABA, suggesting a possible role for ethylene in the very early stages of ripening induction. Altogether, the presented data support a key role for ABA in the regulation of the timing and progress of the complex process of date fruit ripening.

### Endogenous ABA in date fruit ripening

While the role of ABA in the regulation of fleshy fruit ripening is well established, in dates this kind of information is scarce. In the present study, the endogenous ABA levels in date fruit pericarp are shown to rise at the point of fruit green-to-yellow transition, at about 18 WPP, and to consistently increase throughout fruit ripening (Fig. 4). A similar pattern of constant gradual increase in ABA levels, was reported also for fruit ripening in oil palm, strawberry, litchi and other fruit species (Wang *et al*., 2007; Tranbarger *et al*., 2011; Symones *et al*., 2012; Yeap *et al*., 2017*a*). An alternative pattern, in which a transient increase in ABA levels at very early stages of fruit maturation triggers the onset of fruit ripening-related processes, was identified in various fleshy fruit species, including blueberries, grapes, mango, tomato, cucumber, persimmon, and others (Wheeler *et al*., 2009; Zaharah *et al*., 2012; Karppinen *et al*., 2013; Leng *et al*., 2014). In these species, an ethylene peak often accompanies the ABA peak suggesting that both hormones participate in triggering fruit ripening. The pattern of a continuous buildup of endogenous ABA levels, as shown here for date fruit, suggests a relentless and likely multi-layered rather than a transient role for ABA in the regulation of fruit maturation and ripening. Additionally, in oil palm, three ABA-responsive transcription factors were shown to be activated by WRINKELD 1, a master regulator that activates oil synthesis-associated genes, as part of the process of oil palm fruit ripening (Yeap *et al*., 2017*b*). Altogether, the pattern of endogenous-ABA accumulation during the natural process of date fruit development suggests a complex role for ABA in the regulation of date fruit ripening.

### Exogenous ABA applications enhance date fruit ripening

To further characterize the role of ABA in triggering fruit pericarp maturation, date palm samples were exposed to single or repeated exogenous ABA application. The stage at which the hormones were applied had an immense impact on the results obtained. When hormone treatment was applied at 14 WPP, a synergetic effect of the combined ABA and ethylene treatment was observed on fruit pericarp color change (green-to-yellow; Fig. S7), supporting a role for both hormones in early induction of date fruit ripening. In contrast, when applied later in fruit development (16 and 20 WPP), this synergistic hormonal effect was not detected (Figs. S9, S10). Similar synergy between ABA and ethylene was identified in several studies on fleshy fruit ripening in either climacteric or non-climacteric fruits (Zhang, 2014). In litchi, defined as a non-climacteric fruit, ABA did not enhance fruit maturation when applied alone; however, when combined with ethylene, it enhanced both chlorophyll degradation and anthocyanin accumulation, supportive of enhanced fruit ripening (Wang *et al*., 2007). A reciprocity/cross-talk between ethylene and ABA was also reported for grape vine, tomato, fig and persimmon (Zhang *et al*., 2009; Sun *et al*., 2010; Zhao *et al*., 2012; Lama *et al*., 2019). In date palms, a small peak of ethylene production was previously reported at about seed maturation (Serrano *et al*., 2001); yet, so far, this peak was not assigned function in the regulation of fruit ripening. The possibility that a transient peak in ethylene production emerges from the seed-triggering ABA production in the fruit pericarp and thereby induces the process of fruit ripening, requires further research.

As fruit development proceeded, ABA alone was sufficient to enhance date fruit ripening, as indicated by both chlorophyll degradation and sugar accumulation (Figs. 5, 6 and S8). Similar results were reported for other crops. For instance, in mango, which is a climacteric fruit, exogenous ABA treatment promoted fruit pericarp coloration and softening compared to control fruit, while an opposite effect was observed when ABA synthesis was inhibited by nordihydroguaiaretic acid (NDGA) treatment (Zaharah *et al*., 2012). In this context, it is noteworthy that in our experimental setting neither Fluridon, inhibiting the carotenoid pathway, nor NDGA inhibited date fruit ripening (data not shown). In peach, also a climacteric fruit, exogenous ABA application facilitated sugar accumulation (Kobashi *et al*., 2001). Notably, in the current study, single ABA application displayed a short-term effect on the ripening process and hence, multiple applications were required to achieve durable enhancement of fruit ripening and advance fruit harvest. An exception was noted for date palm samples treated at 20 WPP, which showed no measureable effect of ABA on fruit ripening (Fig. S10), suggesting that ABA is sufficient to facilitate date fruit ripening when applied at the appropriate time during fruit development. Taken together, before practical ABA application is considered, it will be essential to determine the optimal treatment schedule and to improve methodologies aiming to enhance the penetration of the active substance into the fruit and to ensure sufficient duration of its impact.

### The final stage in fruit ripening is associated with disruption of xylem-mediated water transport

Due to the harsh arid environment in which dates grow, water dynamics during date fruit development and ripening has been the focus of many studies (Aldrich *et al*., 1946; Rygg, 1946; Gribaa *et al*., 2013; Garcia-Maquilon *et al*., 2017). The modifications in date fruit water status throughout its development are unique; from the initial phase of fruit growth, water levels rise constantly until they reach very sharp maxima as the fruit reaches its final size. From that point and until harvest, fruit WC continuously declines(Bernstein, 2004; Chao and Krueger, 2007; Lobo *et al*., 2013), in parallel to the gradual increase in fruit pericarp ABA levels (Figs. 1, 2 and 4). The relationship between water stress and ABA levels is well studied in a number of fleshy fruit crops (González and Iusem, 2014). A key component identified in that relationship is the water stress-inducible ABA STRESS RIPENING (ASR) transcription factor, which has been well-linked to ripening initiation in crops such as grape, tomato, strawberry, banana and others (Çakir et al., 2003; Golan et al., 2014; Jia et al., 2016; Gupta et al., 2006). ABA also regulates the function of various aquaporins, which mediate water transport between cells and tissues (Fang *et al*., 2019), including in developing fruit (Breia *et al*., 2020). ABA has been also recently implicated in the regulation of the fruit cuticle deposition in orange (Wang *et al*., 2016) and in cucumber (Wang *et al*., 2015) which adds additional aspect to the way it might regulate fruit water status.

After water loss initiation, the next interesting point in water kinetics is the rapid decline in fruit WC just before fruit color transition from yellow to brown. While, xylem-mediated water flux into the fruit was maintained until 20 WPP, at 21-22 WPP, it stopped abruptly just prior to the yellow-to-brown transition (Figs. 3, S4 and S5). Modifications in the vascular tissue during fruit development and ripening were recently identified and thoroughly investigated in grapes. At véraison, the onset of fruit ripening, water flux into the fruit gradually shifts from the xylem to the phloem (Matthews and Shackel, 2005; Choat *et al*., 2009; Keller *et al*., 2015). A more recent study suggested a scenario, in which a discharge of surplus phloem water via berry transpiration and/or xylem backflow to the pedicel, may be necessary to facilitate natural grape ripening as well as sugar accumulation (Zhang and Keller, 2017). Interestingly, a partial fracture of the cluster’s stem, which considerably decreased the water flow to the fruit, was shown to accelerate the ripening of ‘Medjool’ date fruit, when executed at the yellow-to-brown transition, also known as the ‘Khalal-to-Rutab’ shift, suggesting that severe water deficit can trigger the final stage of ripening in mature dates (Bernstein, 2004). Considering the above, we hypothesize that water loss starting at ‘color break’ might elicit a stress signal that induces a series of events facilitating fruit ripening, specifically the last phase of the yellow-to-brown transition.

### Is ABA a multi-tasking hormone?

Soluble sugar levels in ripe fleshy fruits display large variations across species (Coombe, 1976), with very low °Brix (∼ 4) in tomato (Wills and Ku, 2002), moderate (∼11 °Brix) in peach (Bregoli *et al*., 2002), and high (∼24 °Brix) in wine grapes (Bregoli *et al*., 2002; Wills and Ku, 2002; Bondada *et al*., 2017). Date palm fruit are unique in this sense, reaching 44–50 °Brix at full maturity, as the fruit turns from yellow-to-brown (Fig. 1C), and 70 – 80 °Brix at the ripe-dry ready-to-harvest stage, also known as ‘Tamr’ (Chao and Krueger, 2007; Lobo *et al*., 2014). Such high °Brix levels are found only in grape raisins (Aung *et al*., 2002). In both species, the upsurge in fruit sugar concentration is strongly correlated with a significant decline in the fruit WC (Fig. 1C; Aung *et al*., 2002). Within the well-established pivotal and comprehensive role of ABA in the regulation of non-climacteric fruit ripening (Pilati *et al*., 2017; Forlani *et al*., 2019; Fuentes *et al*., 2019; Bai *et al*., 2021), of particular interest is its capacity to integrate changes in fruit sugar metabolism and water status as part of fruit maturation and ripening. Solid evidence suggest a regulatory role for sucrose, in coordination with ABA and other hormones, in the regulation of fruit ripening in grape, strawberry, tomato, apple and peach (Falchi *et al*., 2013; Jia *et al*., 2016, 2017; Ma *et al*., 2017; Olivares *et al*., 2017). In non-climacteric fruit, some evidence suggest a regulatory role for sucrose transporters (*SUT* genes), through which sugars and ABA interact to induce and govern sucrose accumulation and fruit ripening (Jia *et al*., 2013). Additionally, *ASR* gene expression was upregulated by both sucrose and ABA and even more by their combination, which subsequently promoted fruit ripening, whereas RNA interference delayed it (Jia *et al*., 2016*b*). Recently, a study focusing on elucidating ABA signaling pathway in dates identified four *PYRABACTIN-LIKE 8* (*PYL8*) genes, among the 12 genes encoding for the ABA receptor that are expressed during date fruit development. Among these, *PYL8-LIKE 27* was the most highly expressed at the yellow-to-brown transition (Fig. S2; Garcia-Maquilon et al., 2021). The identity of the signal that regulates the timing of the natural increase in fruit pericarp ABA levels and how it is related to the onset of fruit ripening remains to be determined. Various hypotheses can be raised; i) a naturally induced water loss triggers a continuous stress signal, which leads to ABA production and consequently facilitates date fruit maturation and ripening, ii) cross-talk with other hormones that affect ABA metabolism, and last, iii) sucrose levels may regulate ABA accumulation.

### Practical implementations

The observations presented here may have significant practical implications. The date palm industry in Pakistan and India suffers from heavy summer monsoons occurring just before fruit ripening (Abul-Soad *et al*., 2015; Baidiyavadra *et al*., 2019). In northern Africa, early autumn rains and suboptimal temperature often overlap with the date ripening season (Musa, 2001; Awad, 2007; Yahia *et al*., 2013; Jameel M. Al-Khayri *et al*., 2015). In both cases, significant yield proportions are lost. Substantial variability in fruit ripening within a cluster, tree, and orchard cause considerable losses due to unripe or overripe unmarketable fruit. Within-cluster variability also necessitates many rounds of harvest, which is costly and limits date tree orchard size (Musa, 2001; Awad, 2007; Yahia *et al*., 2013; Jameel M. Al-Khayri *et al*., 2015). The ability to hasten date fruit development and regulate the timing of fruit harvest may facilitate expansion of date fruit orchards and cultivation area to habitats with sub-optimal growth conditions.

## Conclusions

Quite similar to véraison in grapes, seed maturation in date palm apprises the onset of fruit maturation and ripening processes in the pericarp, including color change and sugar accumulation. Endogenous ABA gradually rises from the green-to-yellow transition in fruit pericarp, which is equivalent to véraison in grapes, until full ripening. Results of exogenous ABA applications directly associate this hormone with the ripening processes, opening opportunities for practical influence on fruit harvesting time and fruit quality at harvest. Further research is yet required in order to learn what regulates the timing and level of ABA accumulation in fruit pericarp, how it affects the onset of fruit ripening and how it regulates downstream metabolic processes affecting date fruit quality.

## Materials and methods

### Plant material

All experiments were conducted on date trees of the cv. ‘Medjool’ at Grofit orchard, located in the Rift Valley at South Arava district (29° 56′ 27.02″ N, 35° 3′ 52.61″ E, 118 m above sea level), Israel. The experiments were conducted over three growing seasons (2017, 2018 and 2020), on 6-year-old tissue-culture-generated trees, starting from the second year of commercial harvesting. The soil was sandy, composed of 95% sand and 5% silt. In 2017, pollination was carried out with diluted pollen (25% pollen, 75% talc powder) for fruit load control. In 2018 and 2020, the pollen dilution was different (20% pollen, 80% talc powder). In all years, female inflorescences were trimmed as a fruit load management tool.

#### Monitoring of fruit development and ripening over the growing season

Seasonal observations and monitoring of fruit development were carried out from pollination to the beginning of commercial harvest during the years of 2017 and 2018. All time points are presented as weeks post-pollination (WPP). The seasonal observation of date fruit development was conducted in a randomized block design, with 6 single-tree blocks. Four clusters per tree were chosen at anthesis and used for strands and fruit sampling. Eventually, each cluster carried approximately 200 fruits.

#### Pre-harvest single-dose treatment with plant hormones

During the season of 2017, the hormone treatment experiment was laid out in randomized block design. Six trees were chosen at anthesis. One cluster per tree (six trees in total) was used, in which 3-5 strands (with a minimum of 18 fruit per treatment) were used for each treatment. For hormone treatments, 3-5 strands were dipped in plastic bags filled with hormone solution for 15 s, while avoiding exposure of other parts of the tree to the solution. The treatments included untreated control, water control, 0.1% Triton X-100 (Sigma-Aldrich), 6 g L^-1^ (containing 20% S-ABA as the active component) ProTone™ SG (Valent BioSciences) + Triton X-100 (0.1%), 2.1 ml L^-1^ Ethrel™ (Bayer) + Triton X-100 (0.1%) and a combined treatment of 6 g L^-1^ ProTone™ + 2.1 ml L^-1^ Ethrel™ + Triton X-100 (0.1%).

#### Pre-harvest multiple-dose treatment with plant hormones

The 2020 experiments testing pre-harvest exogenous ABA application were extended to whole clusters. Comparable trees in the Grofit orchard were selected. Six uniform clusters from each tree (pollinated in the same week, and each carrying 300 fruits, on average) were chosen, and each one was designated to a different treatment. Thus, the experiment was performed in a randomized block design with 10 trees (each functions as a block) and six treatments per tree. The treatments included untreated control, water control, 0.1% Triton X-100 (Sigma-Aldrich), or 3, 6, or 9 g L^-1^ (containing 20% S-ABA as the active ingredient) ProTone™ SG (Valent BioSciences) + Triton X-100 (0.1%). During application, each cluster was insulated using a plastic conus, to avoid undesired spray to neighboring treated clusters. Clusters were sprayed to drainage to ensure full coverage. Treatments were applied at dawn, at the daily peak of relative humidity and minimum wind, to maximize retention of the spray on the fruit surface. Applications were repeated 4 times, starting in 17 WPP, once every 3 days.

One week after the first application, 3 representative fruits of each cluster were sampled and used for color and Brix determinations. Harvest was carried out in 5 rounds (once a week), upon occurrence of earliest ripening symptoms (one week prior to the start of commercial harvest in the plot). Each cluster was carefully shaken, and the fallen fruit were collected from its net-bag. All fruit from each cluster were counted and sorted according to their ripening stage, as follows: premature dehydration (PMD); yellow-ripe (‘Khalal’); partially-brown; hydrated-brown (‘Rutab’); partially-dry brown (hydrated ‘Tamr’); and, dry brown (dry ‘Tamr’; Figs. 7A and S12). This classification was supported by fruit TSS, which steadily increased with the rise in ripening class (Fig. 7B). All fruits were weighed and the yield was calculated (7B).

### Fruit diameter and weight measurements

Fruit dimensions were measured from top to bottom (calyx to stigma trace; length) and along the mid-fruit equator (diameter) using a high-precision digital caliper (MRC, MT-110051). Fresh and dry weight of the fruit and seed were measured using a precision balance (LE Series, Sartorius or Boeco, BPS 41 Plus). For dry weight and WC calculation, samples were dehydrated using an air-vacuumed oven (Zenith lab, DZF-6050 or WTC Binder, VD-53) for a minimum of 48 h, at 70 °C.

### Total soluble solids measurements

For destructive TSS (Brix units) measurement, fruit pericarp was scraped from top to bottom (from the proximal to the distal side of the fruit). For TSS values <32, measurements were taken using a digital refractometer PR-32α (Atago Co. Ltd.), while for TSS values >32, measurements were taken using an optical refractometer MOD. 103, 0-80 BRIX (Giorgio Bormac S.R.L.).

### Fruit color measurements

Relative fluorescence of detached fruit was measured using a hand-held fluorimeter to estimate chlorophyll levels [Multiplex Research, Force-A; as previously described by **(Bahar *et al***., **2012)**]. The chlorophyll index representing fruit pericarp chlorophyll levels is the calculated SFR_G (simple fluorescence ratio – green excitation) value. Spectral color, using the L*, a*, b* color space coordinates, was measured using either an i1 Pro Rev E Chroma meter (x-rite, PANTON® during the 2017 season) or a CR-400 Chroma meter (Konica Minolta, Inc. during the 2018 and 2020 seasons).

### Histology

Whole fruit and pericarp surfaces were examined using a binocular dissection scope (Nikon, SMZ 1270) and imaged with an attached camera (NIKON, DS-Ri2). Samples were stained with the water soluble Safranin-O dye (Sigma-Aldrich) to examine xylem transport towards the fruit. Six strands were sampled at each time point along the course of fruit development. Three of the strands were placed for 4 h in 5 ml 1% Safranin-O solution and three were placed in 5 ml deionized water as a control. To monitor Safranin-O transport along the xylem and into the fruit pericarp tissue, a longitudinal section of the fruit (through the attached strand segment and at the center of the calyx to the distal end, the stigma trace) and a cross-section (at the middle between the calyx and the distal end) were generated and photographed using zoom stereomicroscope (SMZ1270, Nikon) equipped with a camera (DS-Ri2, Nikon). Fresh lemon juice was applied to avoid oxidation and pericarp browning as a result of the cut.

### ABA quantification

Pericarp tissue (Grofit, 2017 season) was collected at different stages in fruit development (12, 14, 16, 18, 20, 22 and 24 WPP) and frozen in liquid nitrogen. The tissue was ground to a fine powder using a mortar and pestle, after which, 200 mg were weighed and placed in a 2 ml Eppendorf tube containing 1 ml extraction solvent (ES; 80% methanol: 19% water: 1% acetic acid) supplemented with 20 ng of ABA isotope standard (Olchemim). The tube was incubated on Vortex for 60 min at 4 °C and the supernatant was collected following centrifugation. The pellet was washed twice with 0.5 ml ES. The collected supernatant was concentrated with a speed-vac concentrator (5301, Eppendorf), at 30 °C. The samples were then dissolved in 200 μL solution of 50% methanol. Following centrifugation, the supernatant was filtered through a 0.22 μm PVDF syringe filter and then subjected to LC-MS analysis conducted using a UPLC-Triple Quadrupole-MS (Waters Xevo TQ-MS). Separation was performed on a Waters Acquity UPLC BEH C18 1.7 μm 2.1×100 mm column with a VanGuard precolumn (BEH C18 1.7 μm 2.1 × 5 mm). Chromatographic and MS parameters were as follows: the mobile phase consisted of water (phase A) and acetonitrile (phase B), both containing 0.1% formic acid in gradient elution mode. The flow rate was 0.3 ml min^-1^, and column temperature was 35 °C. All analyses were performed using the ESI source in positive ion mode at the following settings: capillary voltage 3.1 KV, cone voltage 30 V, solvation temperature 400 °C, solvation gas flow 565 L/h, source temperature 140 °C. Quantification was performed using MRM acquisition by monitoring the 247/187, 247/173 (RT=3.92, dwell time of 78 msec for each transition) for ABA, and 253/206, 253/234 (RT=3.92, dwell time of 78 msec) for d6-ABA (used as internal standard). Acquisition of LC-MS data was performed using the MassLynx V4.1 software (Waters).

### Statistics

The Tukey-Kramer HSD test was preformed via JMP Pro software (Statistical Discovery™ SAS).

## Abbreviations

ABA: abscisic acid;
WC: water content;
WPP: week post-pollination;

## Supplemental material

Additional supporting information can be found in the online version of this article:

Fig. S1. **Schematic presentation of date palm anatomy and terminology**.

Fig. S2. **The main stages of date palm fruit development**.

Fig. S3. **Date palm fruit development during the 2018 season**.

Fig. S4. **Dye-facilitated assessment of vascular water flux from the strand to the developing fruit**.

Fig. S5. **Vascular water flux is disrupted prior to date fruitlet drop**.

Fig. S6. **Vascular water flux arrest from the strand to the developing date fruit**.

Fig. S7. **The effect of exogenous ABA and ethylene applied 14 WPP, on date fruit development**.

Fig. S8. **The continuous affect of exogenous hormonal treatments applied 14 WPP, on date fruit ripening**.

Fig. S9. **The continuous effect of exogenous hormonal treatments applied 16 WPP, on date fruit ripening**.

Fig. S10. **The effect of exogenous ABA and ethylene applied 20 WPP, on date fruit development**.

Fig. S11. **The effect of pre-harvest ABA treatments on major yield parameters**.

Fig. S12. **The effect of ABA treatment on fruit ripening at harvest**.

## Acknowledgments

The financial support for this research was provided by the Israel Chief Scientist of Agriculture, grant number 12-02-0200 and JNF-KKL for the research in Ramat Negev R&D Center. We also thank Agrica CTS, the Agricultural Division, for contributing ProTone™ required for field experiments. We would like to express our gratitude to the farmers Kobi Tubul and Nimrod Ben-Hur from Grofit orchard for their cooperation and to Dr. Yuval Cohen, Dr. Hamutal Borochov-Neori, Dr. Amnon lichter, Avi Sadovski and Racheli Ben-Zvi for fruitful discussions.

